# SIRT3-dependent mitochondrial oxidative stress in sodium fluoride-induced hepatotoxicity and salvage by melatonin

**DOI:** 10.1101/107813

**Authors:** Chao Song, Jiamin Zhao, Jingcheng Zhang, Tingchao Mao, Beibei Fu, Haibo Wu, Yong Zhang

**Author notes:** These authors contributed equally to this study (Chao Song, Jiamin Zhao). **Corresponding author**: Correspondence should be addressed to Haibo Wu or Yong Zhang., (H.W) or (Y.Z).

## Abstract

Oxidative stress induced by fluoride (F) is associated with fluorosis formation, but the underlying molecular mechanism remains unclear. In this study, Melatonin pretreatment suppressed F-induced hepatocyte injury in HepG2 cells. Melatonin increases the activity of superoxide dismutase (SOD2) by enhancing sirtuin 3 (SIRT3)-mediated deacetylation and promotes SOD2 gene expression via SIRT3-regulated DNA-binding activity of forkhead box O3 (FoxO3a), indicating that melatonin markedly enhanced mROS scavenging in F-exposed HepG2 cells. Notably, melatonin activated the peroxisome proliferator-activated receptor gamma coactivator 1α (PGC-1α). PGC-1α interacted with the estrogen-related receptor alpha (ERRα) bound to the SIRT3 promoter, where it functions as a transcription factor to regulate SIRT3 expression. Furthermore, daily injection of melatonin for 30 days inhibited F-induced oxidative stress in mice liver, leading to improvement of liver function. Mechanistic study revealed that the protective effects of melatonin were associated with down-regulation of JNK1/2 phosphorylation in vitro and in vivo. Collectively, our data suggest a novel role of melatonin in preventing F-induced oxidative stress through activation of the SIRT3 pathway.

## Introduction

Environmental fluoride (F) is a toxic reagent that can affect human health in various ways (Taghipour et al., 2016). Small amounts of F can be used for strengthen bones and prevention of dentals, but excessive F exposure causes a variety of pathological changes in different cells and tissues (Fu et al., 2014). The increasing F concentration in the environment combined with the intervention of human activities is cause for great concern (Ameeramja et al., 2016). The liver is the largest internal organ and the main target of F in the body. Epidemiological and clinical data have shown that excessive sodium fluoride (NaF) exposure results in liver damage (Chattopadhyay et al., 2011). As the cellular outcome of mitochondrial dysfunction, oxidative stress is strongly implicated as one of the most important mechanisms contributing to the toxic effects of NaF (Varol and Varol, 2012). Evidence suggests that the liver is highly vulnerable to oxidative stress because of the accumulation of mitochondrial superoxide anion (O_2_^•−^) (Mahaboob Basha and Saumya, 2013).

As the main mitochondrial deacetylase, sirtuin-3 (SIRT3) modulates various proteins to control mitochondrial function and mitochondrial reactive oxygen species (mROS) (Pi et al., 2015; Sundaresan et al., 2009). Mitochondrial manganese superoxide dismutase (SOD2) is the main enzyme responsible for scavenging harmful O_2_^•−^ and is a substrate of SIRT3 (Kim et al., 2016; Miar et al., 2015). The binding of SIRT3 with SOD2 causes the deacetylation and activation of SOD2 (Tao et al., 2010). Moreover, SIRT3 can also interact with forkhead box O3a (FoxO3a) to activate the FoxO3a-dependent antioxidant-encoding gene SOD2 (Padmaja Divya et al., 2015).

Melatonin and its metabolites are powerful antioxidants and free radical scavengers (Ramis et al., 2015; Siu et al., 2006). Furthermore, melatonin modulates mitochondrial function and strengthens its antioxidant defense systems (Dragicevic et al., 2011). These effects are facilitated by its amphiphilic nature, thereby enabling melatonin to penetrate all morphophysiological barriers and enter all subcellular compartments (Venegas et al., 2012). Recent studies have focused on the role of melatonin in the regulation of mROS levels in healthy and disease states (Acuna Castroviejo et al., 2002). However, the mechanism by which melatonin protects against NaF-induced hepatic oxidative injury remains obscure.

The data presented in the current report are the first to indicate that melatonin efficiently protected against NaF-induced mitochondria-derived and O_2_^•−^-dependent oxidative stress *in vivo* and *in vitro*. All these results contribute to the future clinical treatments of F-induced hepatotoxicity.

## Results

### Melatonin attenuated F-induced oxidative injury in HepG2 cells via SIRT3 pathway

Firstly, we explored the effects of melatonin on F-induced hepatotoxicity *in vitro*. As shown in Fig. 1A, F decreased cell viability in a time- and dose-dependent manner in HepG2 cells. However, melatonin pretreatment significantly attenuated the adverse effect of F on cell viability (Fig. 1B). F treatment resulted in significantly elevated levels of MDA, which was attenuated by melatonin (Fig. 1C). Pre-treatment with melatonin followed by F resulted in restoration of GSH (Fig. 1D). As shown in Figs 1E and F, melatonin protected the HepG2 cells against apoptosis. The expression of Bax was increased significantly in the cytosolic fraction but decreased significantly in the mitochondrial fraction in HepG2 cells when treated with F. The Bax/Bcl-2 ratio after F treatment increased in the cytosolic fraction but significantly decreased in the mitochondrial fraction of HepG2 cells. By contrast, melatonin pretreatment inhibited F-induced changes of Bax/Bcl-2 ratio in the cytosolic and mitochondrial fractions.

**FIGURE 1.**
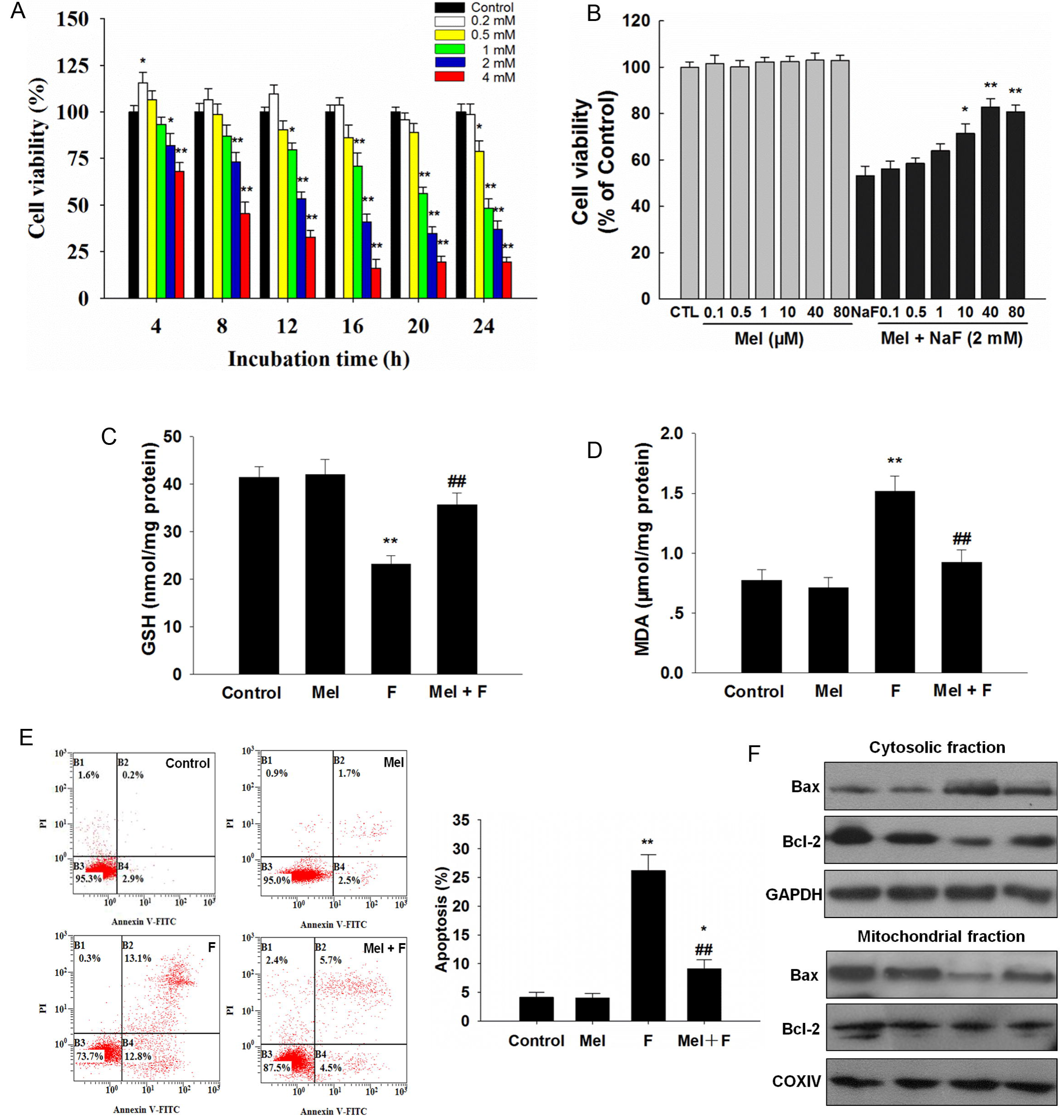
Melatonin inhibited NaF-induced hepatic injury in HepG2 cells. (A) Cells were treated with NaF with different concentrations or for different time intervals, respectively. Cell viability was determined using the CCK-8 assay. (B) Confluent cells were pretreated for 2 h with various concentrations of melatonin. After removing the supernatants, cells were incubated with fresh medium in the presence or absence of NaF (2 mM) for an additional 12 h. Cell viability was determined. (C) GSH level. (D) MDA content. (E) Representative images of flow cytometric analysis by Annexin V-FITC/PI dual staining. (F) The ratio of Bax/Bcl-2 in the cytosolic fraction and mitochondrial fraction. All results are representative of three independent experiments and values are presented as means ± SD. ^*^p < 0.05, ^**^p < 0.01 versus the control group, ^##^p < 0.01 vs. the F group.

As the main mitochondrial deacetylase, sirtuin-3 (SIRT3) modulates various proteins to control oxidative stress response. Melatonin pretreatment significantly recovered the reduced SIRT3 expression (Fig. 2A) and activity (Fig. 2B), which was induced by F. Loss of SIRT3 diminished the effects of melatonin-induced the up-regulated expression and activity of SIRT3. We found that melatonin could protect against F-induced mitochondria-derived O_2_^•−^ elevation (Fig. 2C) and cell viability reduction (Fig. 2D). As shown in Fig. 2E, melatonin significantly blocked the increase in apoptosis induced by F. Furthermore, melatonin treatment inhibited the collapse of MMP induced by F (Fig. 2F). However, these beneficial effects of melatonin were significantly attenuated by SIRT3 siRNA transfection. These data suggest a SIRT3-dependent effect of melatonin on oxidative stress response in hepatic cells exposed to F.

**FIGURE 2.**
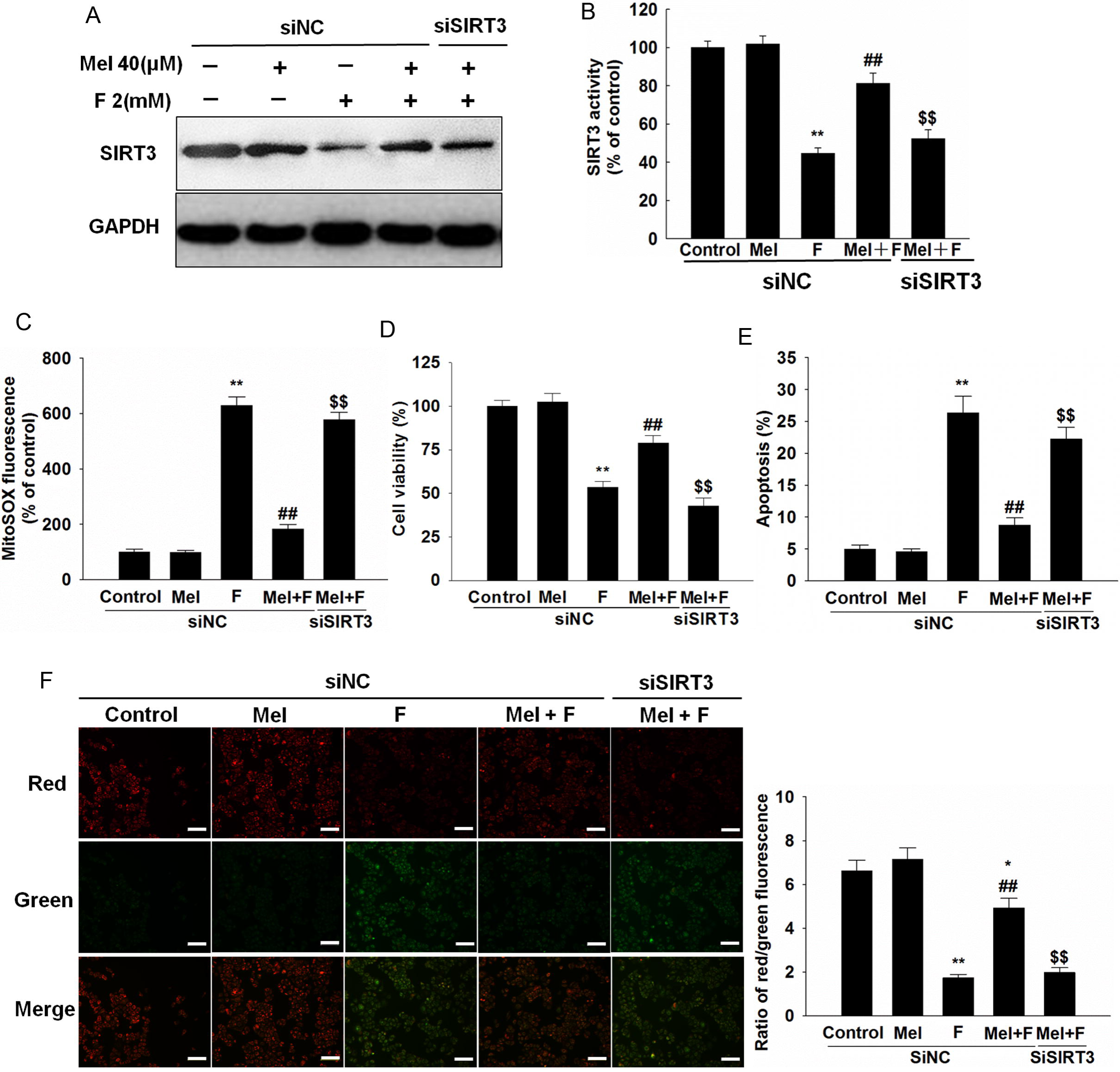
Melatoninprotected against NaF-induced oxidative injury via SIRT3 pathway. Cells were transfected with SIRT3 siRNA. At 24-h post-transfection, cells were pretreated with melatonin (40 μM) for 2 h and then treated with or without NaF of 2 mM for an additional 12 h. SOD2 expression (A) and activity (B), the mitochondrial O_2_•^−^ levels (C), cell viability (D), apoptosis (E), and MMP (F) were determined, respectively. The scale bar is 50 μm. All results are presented as means ± SD of at least three independent experiments. ^*^p < 0.05, ^**^p < 0.01 versus siNC + control group, ^##^p < 0.01 versus the siNC + F group, ^$$^p < 0.01 versus the siNC+ Mel + F group.

Moreover, HepG2 cells were also incubated with F in the presence of Mito-TEMPO (mitochondria-targeted SOD mimetic). Treatment with Mito-TEMPO significantly enhanced SOD2 activity but not SOD2 protein levels (Figs. S1A and B). The F-induced increase in oxidative injury was significantly attenuated in the presence of Mito-TEMPO (Figs. S1c–e).

### Melatonin inhibits mitochondria-derived O_2_^•−^ accumulation via SOD2 upregulation in HepG2 cells

SOD2 is a mitochondrial antioxidant that aids in the elimination of O_2_^•−^. As shown in Figs. 3A and B, loss of SIRT3 diminished the effects of melatonin-induced the up-regulated expression and activity of SOD2. SIRT3-mediated deacetylation of SOD2 and the subsequent regulates its antioxidant activity, we further studied the relationship between the influence of melatonin on SOD2 activity and SIRT3. Coimmunoprecipitation pull-down (Co-IP) assay results indicated that melatonin promoted the binding of SOD2 and SIRT3 in mitochondria under F exposure (Fig. 3C), and caused the decreased acetylation of SOD2 (Fig. 3D). SIRT3 knockdown diminished the effects of melatonin on the acetylation levels of SOD2 (Fig. 3E). Overexpression of SIRT3 significantly rescued F-induced suppression of SIRT3 expression (Fig. 4A) and activity (Fig. 4B). Moreover, overexpression of SIRT3, but not SIRT3^H248Y^ (a catalytic mutant of SIRT3 lacking deacetylase activity), decreased the expression of acetylated-SOD2 and increased SOD2 activity in HepG2 cells exposed to NaF (Figs. 4C and D). The deacetylase-deficient Sirt3 mutant (H248Y) completely eliminated the protective effects of SIRT3 (Figs. 4E and F). These results indicate that the deacetylation of SOD2 induced by melatonin is mediated by SIRT3 and melatonin enhances SOD2 activity through the deacetylation of SIRT3.

**FIGURE 3.**
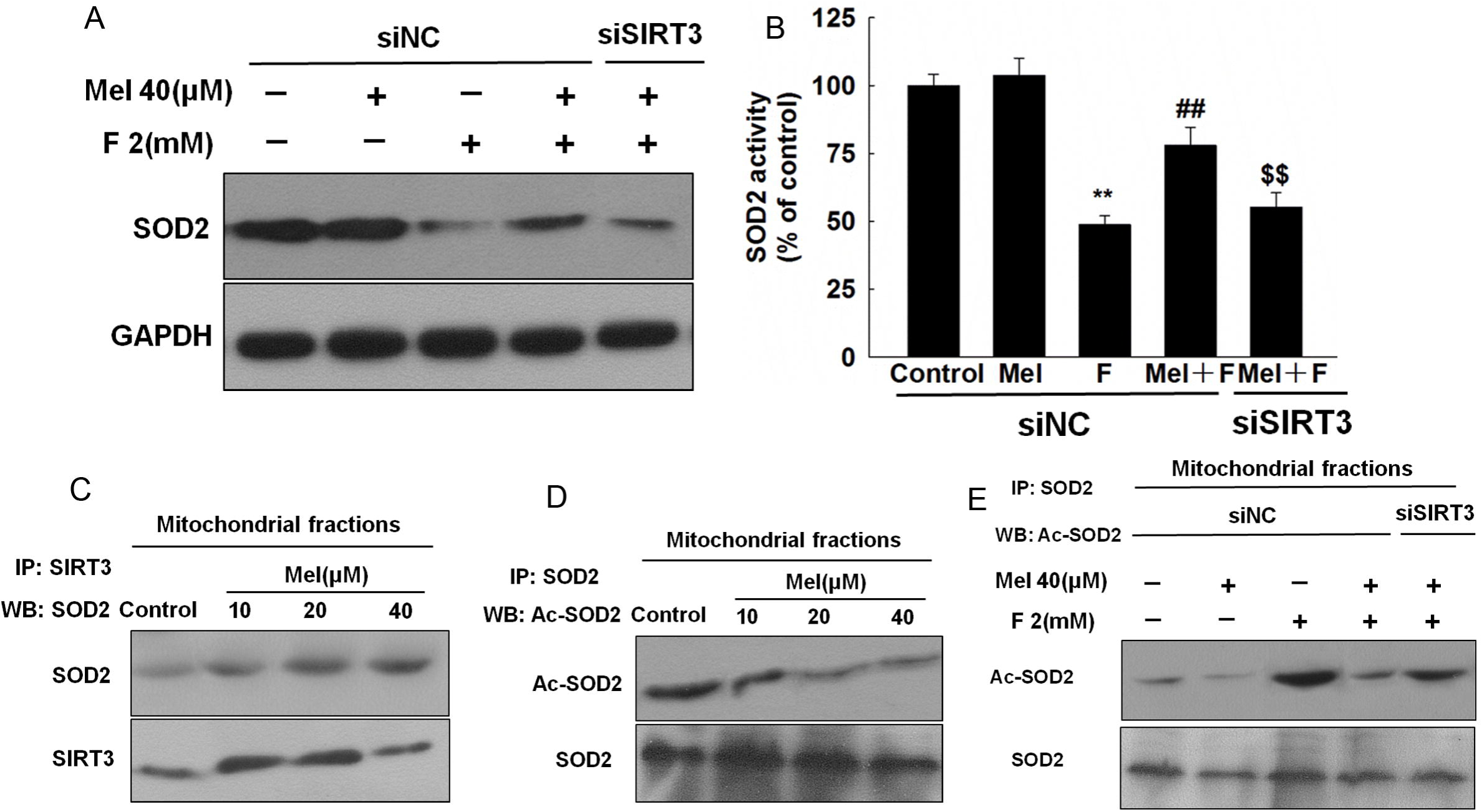
Melatonin increased mROS scavenging by stimulating SIRT3-mediated SOD2 deacetylation. Cells were transfected with SIRT3 siRNA. At 24-h post-transfection, cells were pretreated with melatonin (40 μM) for 2 h and then treated with or without NaF of 2 mM for an additional 12 h. (A) SOD2 expression. (B) SOD2 activity. (C) SOD2 was immunoprecipitated using SIRT3 antibody. (D) acetylated-SOD2 (Ac-SOD2) was immunoprecipitated using SOD2 antibody. (E) Ac-SOD2 was immunoprecipitated in SIRT3-deficient HepG2 cells. All results are representative of three independent experiments. ^**^p < 0.01 versus siNC + control group, ^##^p < 0.01 versus the siNC + F group, ^$$^p < 0.01 versus the siNC+ Mel + F group.

**FIGURE 4.**
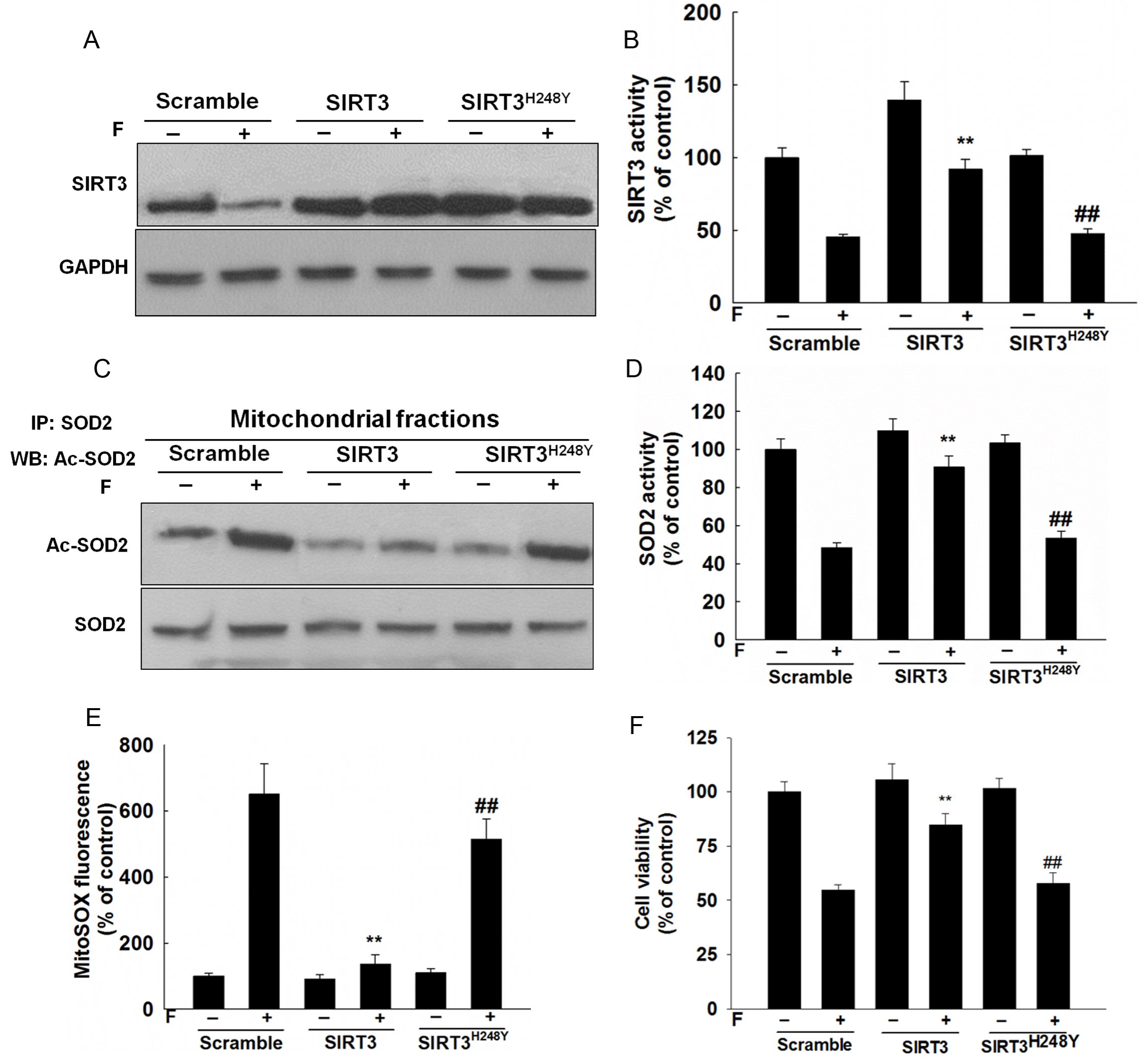
SIRT3 deacetylase deficiency does not affect SOD2 acetylation and oxidative injury in F-treated HepG2 cells. Cells were transfected with SIRT3 expression constructs (WT or H248Y) followed by exposure to NaF (2 mM) for 12 h. (A) SIRT3 expression, (B) SIRT3 activity. (C) Acetylation of SOD2 was determined by immunoprecipitation. (D) SOD2 activity. (E) Mitochondrial-derived O_2_•^−^ production. (F) Cell viability. ^**^p < 0.01 versus scramble + F group, ^##^p < 0.01 versus the SIRT3 + F group.

### Melatonin increased SOD2 expression via the interaction of SIRT3 with FoxO3a

As shown in Fig. 5A, melatonin pretreatment had little influence on the total protein level of FoxO3a in F-treated HepG2 cells. Treatment of HepG2 cells with F caused increased the phosphorylation of FoxO3a at serine 253 (Fig. 5B) and the acetylation of FoxO3a at the K-100 lysine (Fig. 5C). Both events prevented nuclear import, thereby leading to its inactivation. Melatonin pretreatment decreased the phosphorylation at Ser253 and comparably increased FoxO3a deacetylation. Loss of SIRT3 diminished the effects of melatonin on the deacetylation and phosphorylation of FoxO3a. SIRT3 and FoxO3a functionally and physically interact to form a complex that regulates the activity and acetylation status of FoxO3a. Our results showed that melatonin promoted the interaction of FoxO3a with SIRT3 in mitochondria (Fig. 5D). Moreover, the FoxO3a-luciferase reporter gene assay indicated that melatonin increased the transcriptional activity of FoxO3a. When the cells were co-transfected with the pGL-FHRE-luc plasmid and SIRT3 siRNA, the regulative activities of melatonin were abolished (Fig. 5E). The ChIP assay was then performed to investigate the role of FoxO3a in the melatonin-induced SOD2 expression of FoxO3a. In the ChIP assay, melatonin promoted the binding of FoxO3a to the promoter of SOD2 (Fig. 5F) under F exposure.

**FIGURE 5.**
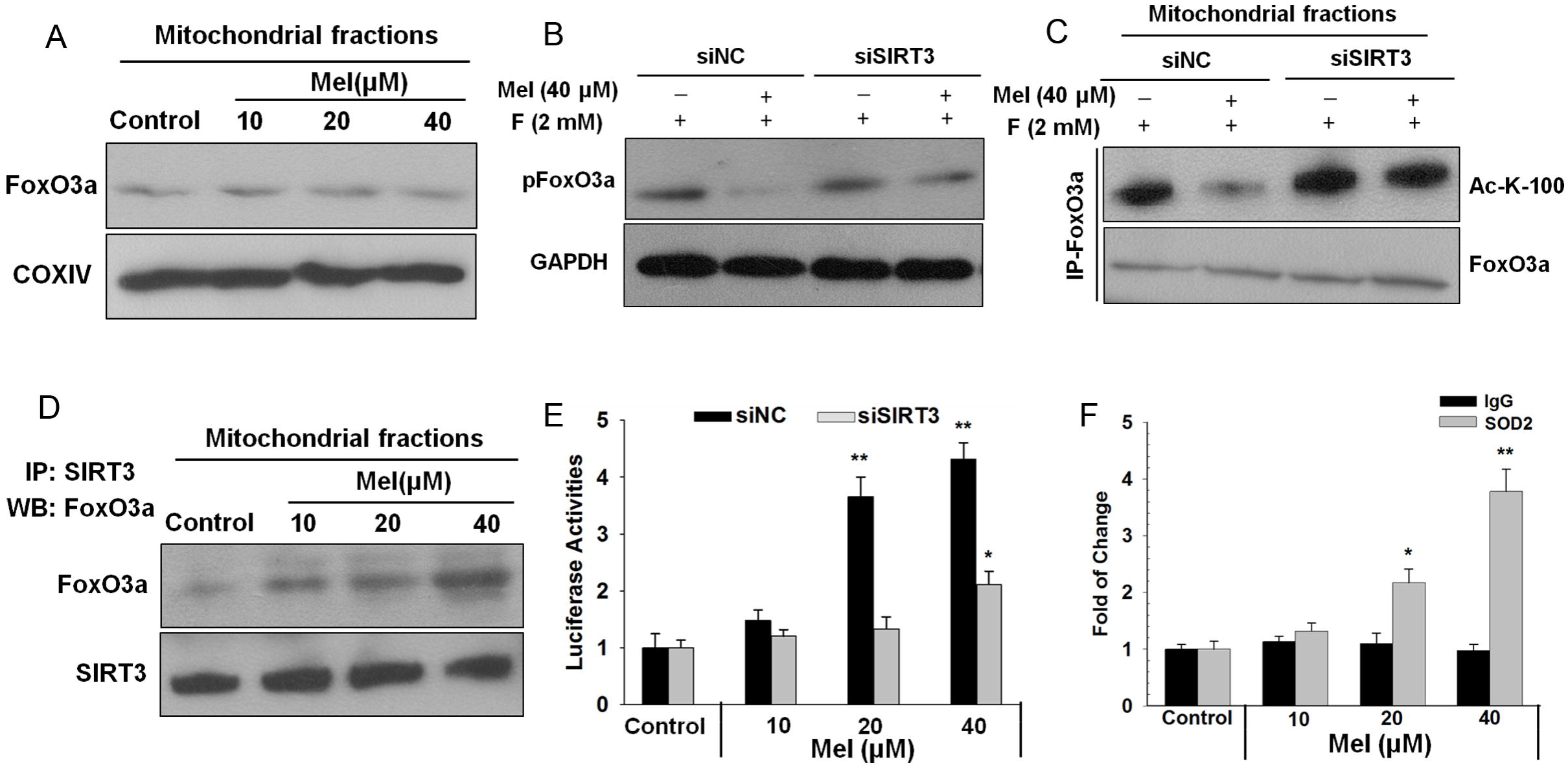
Melatonin regulated the expression of SOD2 through the interaction of SIRT3 with FoxO3a in mitochondria. (A) Mitochondria were isolated after treatment and subjected to western blot analysis for FoxO3a. (B and C) Expressions of p-FoxO3a and FoxO3a acetylation at lysine-100 residue. (D) FoxO3a was immunoprecipitated using a SIRT3 antibody. (E) Cell lysates were harvested for dual luciferase report assays. (F) ChIP analysis was used to examine the binding of FoxO3a to the SOD2 promoter. Data are presented as the mean ± SD of three independent experiments. ^*^P<0.05, ^**^P<0.01 versus the Control group.

The manipulation of SIRT3 expression, but not that of SIRT3^H248Y^, restored activation of FoxO3a by decreasing phosphorylation at Ser253 (Figs. 6A and B) and comparably increasing FoxO3a deacetylation (Fig. 6C). Overexpression of SIRT3 increased the transcriptional activity of FoxO3a (Fig. 6D) and restored the F-induced reduction FoxO3a binding to the gene promoter of SOD2 (Fig. 6E).

**FIGURE 6.**
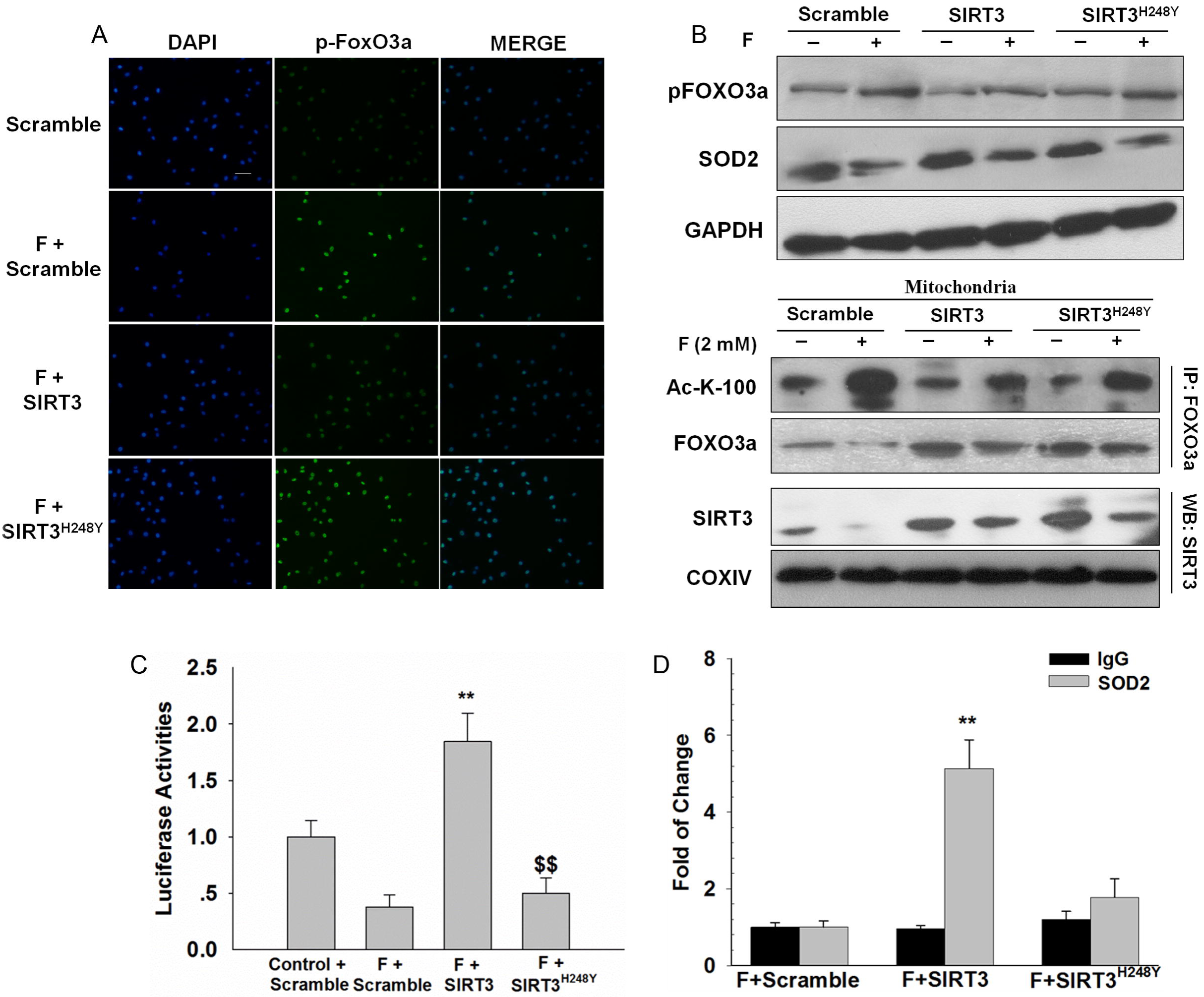
SIRT3 deacetylase deficiency does not affect SOD2 expression and in F-treated HepG2 cells. Cells were transfected with SIRT3 expression constructs (WT or H248Y) followed by exposure to NaF (2 mM) for 12 h. (A and B) The expression of pFoxO3a, FoxO3a, and SOD2. (C) FoxO3a acetylation at lysine-100 residue was examined. (D) Cell lysates were harvested for dual luciferase report assays. (E) Binding of FoxO3a to SOD2 was examined using ChIP assay. The scale bar is 50 μm. Data are presented as the mean ± SD of three independent experiments. ^**^P<0.01 versus the F + Scramble group, ^$$^P<0.01 versus the F + SIRT3 group.

### Melatonin activated the PGC-1α/ERRα-SIRT3 signaling pathway in HepG2 cells

Next, we sought to determine which factor mediates the expression of SIRT3. As shown in Figure. 7A, SIRT3 expression was regulated at the transcription level. SIRT3 was previously reported to be regulated by PGC-1a. So we examined PGC-1a expression with western blotting and found that PGC-1a was changed with a similar change pattern of SIRT3 expression (Fig.2B). PGC-1α siRNA transfection prevented the induction of SIRT3 expression in HepG2 cells, thereby indicating that PGC-1α was required for the activation of melatonin during SIRT3 expression.

**FIGURE 7.**
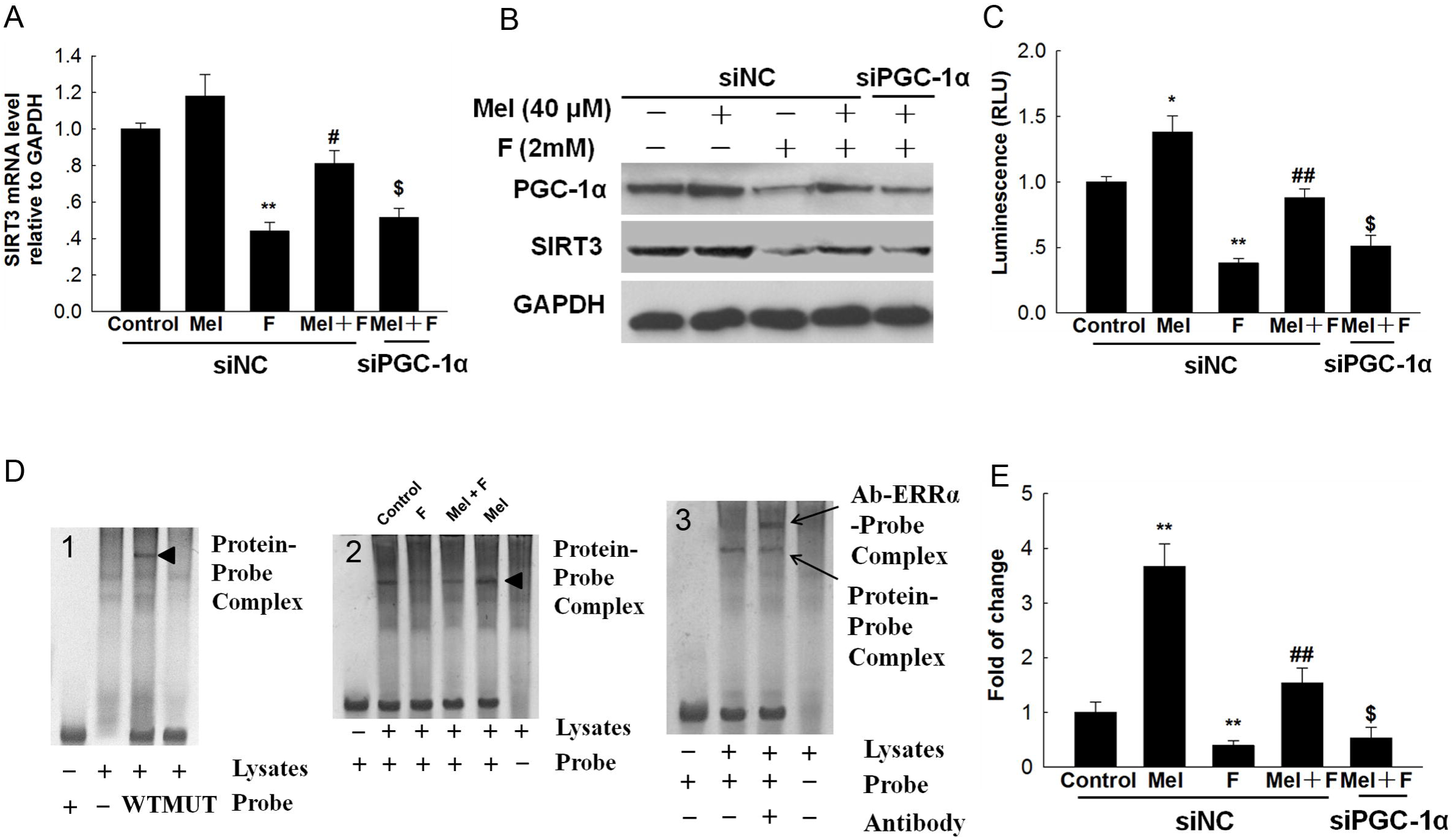
Melatonin activated PGC-1α/ERRα-SIRT3 signaling pathway in HepG2 cells. Cells were transfected with PGC-1α siRNA. At 24-h post-transfection, cells were pretreated with melatonin (40 μM) for 2 h and then treated with or without NaF of 2 mM for an additional 12 h. (A) SIRT3 mRNA level. (B) The expression of PGC-1a and SIRT3. (C) ERRα luciferase reporter plasmid was transfected into HepG2 cells. Luciferase activity was determined. (D) *In vitro* binding of ERRα and SIRT3 promoter was examined by EMSA assay. (E) *In vivo* binding of ERRα and SIRT3 promoter was examined by the ChIP assay. Data are presented as the mean ± SD of three independent experiments. ^*^p < 0.05, ^**^p < 0.01 versus the siNC + control group; ^#^p < 0.05, ^##^p < 0.01 versus the siNC + F group, and ^$^p < 0.05 versus the siNC +Mel + F group. Abbreviation: WT, wild-type; MUT, mutation; Ab, antibody.

Since ERRα acts as the downstream target of PGC-1α and is co-activated by this transcriptional coactivator. Here we found that PGC-1α interacted with ERRα in HepG2 cells (Fig. S2A). Knockdown of ERRα or PGC-1α decreased SIRT3 expression and cotransfection of ERRα siRNA and PGC-1a siRNA could decrease more expression of SIRT3 (Fig. S2B), while overexpression of PGC-1α or ERRα increased SIRT3 expression and cotransfection of PGC-1α and ERRα could increase more expression of SIRT3 (Fig. S2C). These results indicate that PGC-1α and ERRα coordinately regulate SIRT3 expression in HepG2 cells. Furthermore, the luciferase assay was used to determine if SIRT3 mRNA activation by melatonin occurred via PGC-1α-dependent ERRα binding to the SIRT3 promoter. As shown in Fig. 7C, F exposure caused a significant decrease in the luciferase activity as compared with the control group. By contrast, melatonin pretreatment significantly increased the ERRE-mediated SIRT3 transcriptional activity, whereas this effect was diminished by the addition of PGC-1α siRNA.

The EMSA assay was further performed to test the *in vitro* binding of ERRα and SIRT3 fragments. A preliminary experiment was performed to test the binding of probe (Fig. 7D-1). As shown in Fig. 7D-2, a band in the lane loaded with WT probe and lysates was shifted as compared with lysates alone. The protein–probe binding was regulated by F and melatonin. Moreover, a specific super-shift band was detected with the ERRα antibody, thereby indicating that ERRα was bound to the probe (Fig. 7D-3). The results of EMSA also showed that melatonin increased the binding of the exogenous consensus DNA oligonucleotide of SIRT3 with ERRα. To further confirm the EMSA results and verify that ERRα physically occupies the SIRT3 promoter, we performed the ChIP assay. As shown in Fig. 7E, a 3.6-fold enrichment of ERRα was observed.

### Melatonin attenuated F-induced JNK1/2 activation in mice liver

Furthermore, we also investigate whether melatonin could prevent F-induced oxidative stress in mice liver. Significant accumulation of F was observed in F-toxicated mice liver (Fig. S3A). Liver functions were also measured based on changes in the hepatic markers ALT and AST. The results showed that the serum activities of ALT and AST were significantly increased in the F group when compared with the control group (Figs. S3B and C). This result confirmed that F-toxicity model had been successfully established. Melatonin supplementation caused a significant reduction in the accumulated F-content and the serum activities of ALT and AST. Moreover, Melatonin treatment successfully attenuated the F-induced upregulation of O_2_^•−^ and MDA content (Figs. S3D and E). Moreover, GSH level were decreased significantly in F-toxicated mice livers, which was reversed by melatonin (Fig. S3F).

Since apoptosis plays an important role in the pathogenetic mechanisms involved in fluorosis. NaF treatment increased caspase-3 activity (Fig. 8A), an indicator of apoptosis, and decreased the protein levels of Bcl-2 (Fig. 8B), an important anti-apoptotic factor in mice liver. Pretreatment with melatonin attenuated caspase-3 activity and increased Bcl-2 protein expression in NaF-treated mice liver. Activation of mitogen-activated protein kinase (MAPK) has been implicated in F-induced apoptosis and they are sensitive to oxidative stress, Western blot showed that the melatonin significantly reduced the phosphorylation of JNK1/2 in F-exposed mice liver.

**FIGURE 8.**
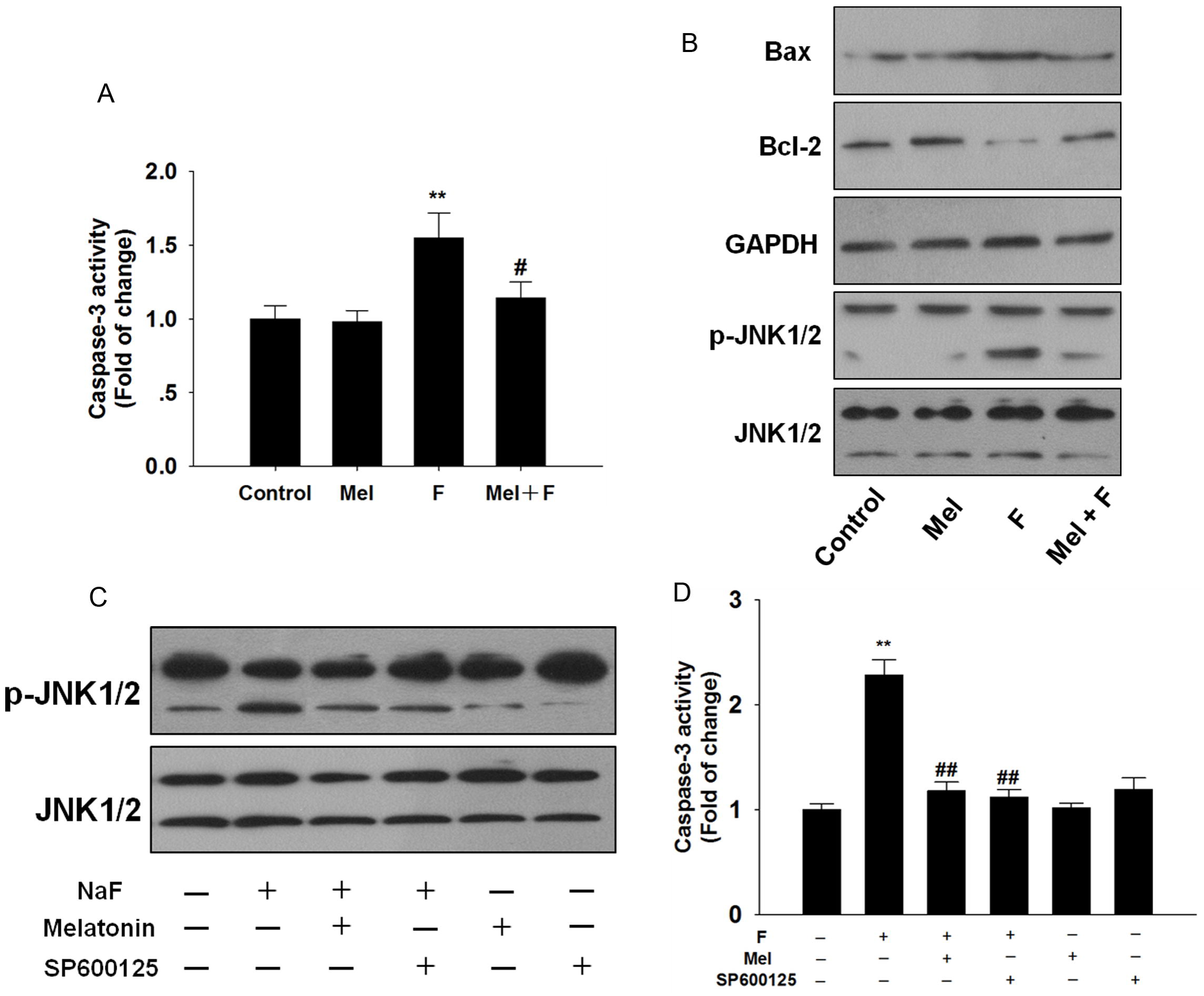
Melatonin inhibited JNK1/2 activation in mice liver. (A) Caspase-3 activity (H). (B) The representative western blot for Bax, Bcl-2, JNK1/2 in mice liver. (C) Immunoblot analysis in HepG2 cells. (D) Caspase-3 activity was determined in HepG2 cells. Data are mean ± SD; n = 6–8. ^*^p < 0.05 or 3 different cultures., ^**^p < 0.01 versus the control group, ^#^p < 0.05, ^##^p < 0.01 versus the F group.

To further address the involvement of JNK1/2, HepG2 cells were pretreated melatonin or SP600125 (a potent JNK1/2 inhibitor). Our results showed that F-induced JNK1/2 activation in HepG2 cells, which was significantly reduced by melatonin or SP600125 (Fig. 8C). In addition, caspase-3 activity were reduced in F-treated HepG2 cells following pretreatment with melatonin or JNK inhibitor (Fig. 8D).

## DISCUSSION

Oxidative stress from excessive mROS plays an important role in the pathogenesis of F-mediated cytotoxicity (Chouhan and Flora, 2008; Gao et al., 2008). SOD2 is crucial for maintaining the mitochondria-derived O_2_^•−^ balance (Li et al., 2004). In the present study, SOD2 expression and activity were significantly reduced in NaF-treated HepG2 cells. Pretreatment with melatonin promotes SOD2 activity and expression, thereby protecting liver cells from mROS induced oxidative stress under excessive F exposure. Moreover, treatment with the mitochondria-targeted antioxidant Mito-TEMPO alleviated cellular oxidative stress and increased cell viability. The maintenance of mitochondria-derived O_2_^•−^ at tolerable levels may be a viable strategy to treat F-induced cellular damage.

Oxidative stress is the cellular outcome of mitochondrial dysfunction. The MMP is a measure of mitochondrial function (Gong et al., 2014). The diminished membrane potential in the present study is consistent with previous findings in fluorosis (Yan et al., 2015). Moreover, mitochondrial dysfunction was further aggravated by the down regulation of SOD2 which is critical for maintaining mitochondrial oxidative balance. As the primary mitochondrial deacetylase, SIRT3 regulates the biological functions directly involved in mitochondrial function and oxidative stress response (Ansari et al., 2016; Qiu et al., 2010). SIRT3 also has a positive role in maintaining the MMP (Zhou et al., 2014). Melatonin treatment ameliorates mitochondrial oxidative stress by scavenging for enhanced mROS and improving the mitochondrial function.

SOD2 is a substrate of SIRT3 in mitochondria; the binding of SIRT3 with SOD2 causes SOD2 deacetylation, thereby enhancing the mitochondrial scavenging capacity (Chen et al., 2011; Qiu et al., 2010). Our current findings indicate that melatonin promotes SOD2 deacetylation in mitochondria, which was suppressed by SIRT3 siRNA transfection. Furthermore, a catalytic mutant of SIRT3 (SIRT3^H248Y^), which lacks deacetylase activity, failed to reverse the F-induced increase in mitochondria-derived O_2_^•−^. Therefore, SIRT3 activates SOD2 via its deacetylase activity to inhibit O_2_^•−^ accumulation.

SIRT3 directly targets SOD2 and promotes SOD2 transcription by FoxO3a activation, thereby protecting cells from cellular oxidative stress (Du et al., 2013; Wei et al., 2015). Nuclear localization of FoxO3a is essential for its transcriptional activity and the transcription of FoxO3a-dependent genes (Calnan and Brunet, 2008; Tseng et al., 2013). In the present study, F treatment increased FoxO3a phosphorylation at Ser253, thereby inactivating FoxO3a and preventing its nuclear import to inactivate HepG2 cells. Melatonin promotes the interaction of FoxO3a with SIRT3 and enhances FoxO3a deacetylation in the mitochondria; thus the translocation of FoxO3a from mitochondria to nucleus is promoted, thereby causing its complex influence on SOD2 transcription, which controls mitochondria-derived O_2_^•−^. Collectively, our findings indicate that melatonin enhances SOD2 expression by promoting SIRT3-regulated FoxO3a transcriptional activity under excessive F exposure.

Melatonin-induced SIRT3 activation plays a pivotal role in protecting against F-induced oxidative stress. The transcriptional coactivator PGC-1α is a crucial regulator in mitochondrial biogenesis, energy generation, and oxidative stress response (Houten and Auwerx, 2004; Millay and Olson, 2013). Furthermore, a current study indicated that PGC-1α stimulated mouse SIRT3 activity in both hepatocytes and muscle cells, indicating that PGC-1α acts as an endogenous regulator of SIRT3 (Park et al., 2012). Recent work highlights the key role of PGC-1α in melatonin-regulated hepatic mitochondrial health. Our study is the first to reveal that melatonin activates SIRT3 mRNA transcription via a PGC-1α-dependent signaling pathway. PGC-1a interacts with the SIRT3 transcription factor ERRα as a transcriptional coactivator for the expression of SIRT3 in F-induced liver cells injury.

Oxidative stress have been demonstrated to activate a variety of signaling pathways, among which ERK1/2 and JNK1/2 have been implicated in F-induced apoptosis (Geng et al., 2014). Here, melatonin pretreatment prevented JNK1/2 activation induced by F in vivo and in vitro. Inhibition of JNK1/2 prevented F-induced apoptosis. Thus, blocking JNK1/2 signaling may represent a potential mechanism underlying the protection of melatonin.

In summary, we obtained evidence for the first time to demonstrate mROS inhibition by melatonin reduces F-induced oxidative stress via SIRT3 upregulation in HepG2 cells. Melatonin enhances SOD2 activity by promoting SIRT3-mediated deacetylation, but also increases SOD2 expression by increasing the transcriptional activity of FoxO3a. Our results further highlight the potential importance of melatonin in SIRT3-mediated mROS homeostasis, thereby illustrating a novel molecular mechanism of melatonin to be explored for future clinical treatment of F-induced hepatotoxicity.

## Materials and methods

### Ethics statement

This study was performed in strict accordance with the guidelines for the care and use of animals of Northwest A&F University. All animal experimental procedures were approved by the Animal Care Commission of the College of Veterinary Medicine, Northwest A&F University.

### Chemicals and reagents

All chemicals and reagents were obtained from Sigma-Aldrich Chemical Co. (St. Louis, MO, USA) unless otherwise stated. Antibodies against Bax, Bcl-2, GAPDH, β-actin, COX IV, SOD2, SIRT3, acetylated-lysine, ERRα, and c-Jun NH2-terminal kinase-1/2 (JNK1/2) were purchased from Cell Signaling Technology (Beverly, MA, USA). Antibodies against FoxO3a, phospho-FoxO3a (Ser 253), and PGC-1α were obtained from Abcam (Cambridge, UK).

### Cell culture and treatment

HepG2 cells were purchased from the American Type Culture Collection (ATCC, Manassas, USA). The HepG2 cells were cultured in Dulbecco’s Modified Eagle’s Medium (DMEM, Gibco, USA), which was supplemented with 10 % heat-inactivated fetal bovine serum (FBS, Gibco) in a 5 % CO_2_ humidified atmosphere at 37 °C. Cells were pretreated with 40 μM melatonin for 2 h, washed, and treated with or without NaF (2 mM) for an additional 12 h. Experimental protocol are described in more details in Supporting information.

### Cell viability

Cell viability was analyzed using Cell Counting Kit-8 (Beyotime, Jiangsu, China). The absorbance was obtained with a microplate reader (Epoch, BioTek, Luzern, Switzerland) set at a wavelength of 450 nm.

### Assay of biochemical makers of oxidative stress

The concentrations of malondialdehyde (MDA) and glutathione (GSH), and the activity of SOD2 were assayed by commercial assay kits purchased from Nanjing Jiancheng Bioengineering Institute (Nanjing, Jiangsu, China).

### Apoptosis analysis

Cell apoptosis was detected with the Annexin V–fluorescein isothiocyanate (FITC) Apoptosis Detection kit (Beyotime) and analyzed on the BD LSR II flow cytometry system (Becton Dickinson, Franklin Lakes, NJ, USA).

### Isolation of cytosolic and mitochondrial fractions

Mitochondrial fractions were immediately extracted with the Cytosolic and Mitochondria Isolation Kit (Beyotime). Protein concentrations were determined with the BCA Protein Assay Kit (Pierce Biotech, Rockford, IL, USA).

### Immunoblot analysis

Immunoblot analysis was performed as described in the protocols provided by the primary antibody suppliers.

### SIRT3 activity

Protein was extracted with a mild lysis buffer (50□mM Tris-HCl, pH 8; 125□mM NaCl; 1□mM DTT; 5□mM MgCl_2_; 1□mM EDTA; 10 % glycerol; 0.1% NP-40). SIRT3 activity was determined with the CycLex SIRT3 Deacetylase Fluorometric Assay Kit according to the manufacturer’s instructions (MBL International Corp. Tokyo, Japan). The fluorescence intensity was monitored at excitation and emission wavelengths of 355 and 460 nm, respectively.

### Mitochondrial O_2_^•−^ assessment

Mitochondrial O_2_^•−^ generation was assessed in HepG2 cells by 10 μM MitoSOX (Molecular Probes, CA, USA) for 20 min at 37 °C. The fluorescence intensity was measured with an Infinite™ M200 Microplate Reader (Tecan, Mannedorf, Switzerland) at excitation and emission wavelengths of 492 and 595 nm, respectively.

### RNA interference

The siRNA targeting SIRT3 and PGC-1α were purchased from Santa Cruz Biotechnology (Santa Cruz, CA, USA). Cells were transfected with the non-targeted control siRNA to target small interfering RNAs for 6 h according to the manufacture's protocol. At 24 h after transfection, cells were harvested for further experiments.

### Mitochondrial membrane potential (MMP)

MMP was detected with the fluorescent, lipophilic dye, JC-1 (BioVision, Milpitas, CA, USA) as previously described (Ye et al., 2011).

### Immunoprecipitation (IP)

IP was conducted according to previously described methods (Lai et al., 2013) with a few modifications. Lysates were clarified by centrifugation at 14,000×g for 15 min and adjusted to the same protein concentration with the respective lysis buffer for IP. Briefly, protein extracts were incubated overnight at 4 °C with the anti-SIRT3, SOD2 or FoxO3a antibody before fresh protein A/G-conjugated beads (Santa Cruz) were added and rotated overnight at 4 °C. Finally, the beads were washed thrice with the same lysis buffer, eluted with the sample loading buffer, and subjected to immunoblot analysis.

### Plasmids and transfection

SIRT3 cDNA was cloned from HepG2 cells and inserted into the *Eco*RI/*Xho*I site of the vector pIRES-hrGFP-1a. The mutant of SIRT3 (H248Y) was generated with the Quick Change Site-directed Mutagenesis Kit (Stratagene, Santa Clara, CA, USA). Cells were washed after 24 h of transfection and processed for further studies. All the primers used for plasmids construction are listed in Supporting Information Table S1.

### Immunofluorescence staining

Cells were fixed with 4% formaldehyde, permeabilized with 0.5% Triton X-100, washed with PBS, blocked for 1 h with 10 % bovine serum albumin, and incubated with rabbit monoclonal anti-pFoxO3a (S253) overnight at 4 °C. The cells were then washed and incubated with FITC- conjugated secondary antibodies (Santa Cruz) for 1 h at room temperature, and nuclei were revealed with DAPI (10 mg/ml; Sigma–Aldrich Co., St. Louis, MO). The stained cells were observed by fluorescence microscopy (Nikon, Tokyo, Japan).

### Luciferase reporter assay

Luciferase measurements were performed with the dual luciferase reporter (DLR) Assay System (Promega, Madison, WI, USA). pGL-FHRE-Luc (reporter plasmid for FoxO3a) and pGL-ERRE-Luc (reporter plasmid for ERRα) plasmids were obtained from Addgene (Cambridge, MA, USA). Briefly, cells were transfected with 2 μg of reporter plasmid/well and 0.1 μg of *Renilla* luciferase plasmid pRL-SV40 (Promega) was co-transfected as an internal control. Data were collected with a VICTOR X5 Multilabel Plate Reader (PerkinElmer).

### Quantitative Real-time PCR (qPCR) analysis

Total RNA was isolated with the TRIzol Reagent (Invitrogen), which was reverse transcribed to cDNA with the SYBR PrimeScript RT-PCR Kit (Takara BIO Inc., Japan). The gene-specific primers used are listed in Supporting Information Table S2. Results were normalized to levels of GAPDH mRNA and expressed as the fold change (2^−ΔΔCt^).

### Chromatin immunoprecipitation assay (ChIP)

A ChIP assay was performed with the Pierce Agarose ChIP Kit as previously described (Wu et al., 2014). Briefly, cells were cross-linked with formaldehyde for 15 min at room temperature followed by glycine treatment to stop the cross-linking. Genomic DNA was isolated and sheared by ultrasonic waves and 10 % of the supernatant was regarded as input. Antibodies against FoxO3a or ERRα were used for IP. The ChIP enrichment was determined with an ABI StepOnePlus PCR system (Applied Biosystems). Primer sequences used for ChIP-qPCR are listed in Supporting Information Table S3.

### Electrophoretic mobility shift assay (EMSA)

The EMSA assay was strictly performed with an Electrophoretic Mobility Shift Assay Kit (Molecular Probes, Invitrogen) according to the manufacturer’s recommendations. Primer sequences used for EMSA are listed in Supporting Information Table S4.

### Animal studies

A total of 40 two-mo-old Kunming mice were purchased from the experimental animal center of the Fourth Military Medical University. The mice were kept in standard animal housing at 22 ± 2 °C with ventilation and hygienic conditions, as well as free access to food and water.

Animals were randomly divided into four groups of 10 each. Group 1 (Control): Mice was provided distilled water and received daily injection vehicle for 30 days. Group 2 (Mel): Mice were received daily injection of melatonin alone (5 mg/kg/day, i.p.) for 30 days (San-Miguel et al., 2015). Group 3 (F): Mice were supplied with 120 mg/L NaF in deionized water and received daily injection vehicle for 30 days according to our previous study (Fu et al., 2014). Group 4 (Mel + F): Mice were administrated with 120 mg/L NaF in drinking water and received daily injection of melatonin for 30 days. After treatment, the F concentration in the liver was estimated with an ion-sensitive electrode, as previously described (Zhou et al., 2013). Values were expressed as μg F per g dry tissue.

The livers were homogenized in nine fold (w/v) cold normal saline using an automatic homogenizer, and centrifuged at 1500×g for 20 min at 4 °C. The supernatant was kept at –80 °C until further analysis.

### Liver function

Liver function was evaluated by measuring serum alanine aminotransferase (ALT) and aspartate aminotransferase (AST) with an automated chemistry analyzer (Olympus AU1000, Olympus, Tokyo, Japan).

### Statistical analysis

Raw data were analyzed with the SPSS 19.0 software (Chicago, IL, USA). Results are expressed as mean ± SD from triplicate parallel experiments unless otherwise indicated. Statistical analyses were performed with one-way ANOVA, followed by post hoc least significant difference tests. Values with P<0.05 were considered statistically significant.

## Acknowledgments

This work was funded by the State Key Program (No. 31530075) of National Natural Science Foundation of China and China Postdoctoral Science Foundation Grant (No. 2016M590978).

### Competing interests

The authors indicate no competing financial interest.

### Author contributions

Chao Song: Conception and design, collection and assembly of data, data analysis and interpretation, manuscript writing

Jiamin Zhao: Collection and assembly of data

Jingcheng Zhang: Collection and assembly of data

Tingchao Mao: Collection and assembly of data

Beibei Fu: Collection and assembly of data

Haibo Wu: Conception and design, collection and assembly of data, data analysis and interpretation

Yong Zhang: Conception and design, data analysis and interpretation, financial support, final approval of manuscript

### Abbreviations

NaF: Sodium fluoride
Mel: Melatonin
mROS: Mitochondrial reactive oxygen species
SOD2: Mitochondrial manganese superoxide dismutase
PGC-1α: Peroxisome proliferator-activated receptor gamma coactivator 1α
ERRα: Estrogen-related receptor alpha
MDA: Malondialdehyde
GSH: Glutathione
ChIP: Chromatin immunoprecipitation assay
EMSA: Electrophoretic mobility shift assay
ALT: Alanine aminotransferase
AST: Aspartate aminotransferase
JNK1/2: d c-Jun NH2-terminal kinase-1/2
WT: wild-type
MUT: mutation
Ab: antibody

## References

Acuna Castroviejo, D., Escames, G., Carazo, A., Leon, J., Khaldy, H. and Reiter, R. J. (2002). Melatonin, mitochondrial homeostasis and mitochondrial-related diseases. Curr Top Med Chem 2, 133–51.

Ameeramja, J., Panneerselvam, L., Govindarajan, V., Jeyachandran, S., Baskaralingam, V. and Perumal, E. (2016). Tamarind seed coat ameliorates fluoride induced cytotoxicity, oxidative stress, mitochondrial dysfunction and apoptosis in A549 cells. J Hazard Mater 301, 554–65.

Ansari, A., Rahman, M. S., Saha, S. K., Saikot, F. K., Deep, A. and Kim, K. H. (2016). Function of the SIRT3 mitochondrial deacetylase in cellular physiology, cancer, and neurodegenerative disease. Aging Cell.

Calnan, D. R. and Brunet, A. (2008). The FoxO code. Oncogene 27, 2276–88.

Chattopadhyay, A., Podder, S., Agarwal, S. and Bhattacharya, S. (2011). Fluoride-induced histopathology and synthesis of stress protein in liver and kidney of mice. Arch Toxicol 85, 327–35.

Chen, Y., Zhang, J., Lin, Y., Lei, Q., Guan, K. L., Zhao, S. and Xiong, Y. (2011). Tumour suppressor SIRT3 deacetylates and activates manganese superoxide dismutase to scavenge ROS. EMBO Rep 12, 534–41.

Chouhan, S. and Flora, S. J. S. (2008). Effects of fluoride on the tissue oxidative stress and apoptosis in rats: Biochemical assays supported by IR spectroscopy data. Toxicology 254, 61–67.

Dragicevic, N., Copes, N., O'Neal-Moffitt, G., Jin, J., Buzzeo, R., Mamcarz, M., Tan, J., Cao, C., Olcese, J. M., Arendash, G. W. et al. (2011). Melatonin treatment restores mitochondrial function in Alzheimer's mice: a mitochondrial protective role of melatonin membrane receptor signaling. J Pineal Res 51, 75–86.

Du, K., Yu, Y., Zhang, D., Luo, W., Huang, H., Chen, J., Gao, J. and Huang, C. (2013). NFkappaB1 (p50) suppresses SOD2 expression by inhibiting FoxO3a transactivation in a miR190/PHLPP1/Akt-dependent axis. Mol Biol Cell 24, 3577–83.

Fu, M., Wu, X., He, J., Zhang, Y. and Hua, S. (2014). Natrium fluoride influences methylation modifications and induces apoptosis in mouse early embryos. Environ Sci Technol 48, 10398–405.

Gao, Q., Liu, Y. J. and Guan, Z. Z. (2008). Oxidative stress might be a mechanism connected with the decreased alpha 7 nicotinic receptor influenced by high-concentration of fluoride in SH-SY5Y neuroblastoma cells. Toxicol In Vitro 22, 837–43.

Geng, Y., Qiu, Y., Liu, X., Chen, X., Ding, Y., Liu, S., Zhao, Y., Gao, R., Wang, Y. and He, J. (2014). Sodium fluoride activates ERK and JNK via induction of oxidative stress to promote apoptosis and impairs ovarian function in rats. J Hazard Mater 272, 75–82.

Gong, S., Peng, Y., Jiang, P., Wang, M., Fan, M., Wang, X., Zhou, H., Li, H., Yan, Q., Huang, T. et al. (2014). A deafness-associated tRNAHis mutation alters the mitochondrial function, ROS production and membrane potential. Nucleic Acids Res 42, 8039–48.

Houten, S. M. and Auwerx, J. (2004). PGC-1alpha: turbocharging mitochondria. Cell 119, 5–7.

Kim, H., Lee, Y. D., Kim, H. J., Lee, Z. H. and Kim, H. H. (2016). SOD2 and Sirt3 Control Osteoclastogenesis by Regulating Mitochondrial ROS. J Bone Miner Res.

Lai, L., Yan, L., Gao, S., Hu, C. L., Ge, H., Davidow, A., Park, M., Bravo, C., Iwatsubo, K., Ishikawa, Y. et al. (2013). Type 5 adenylyl cyclase increases oxidative stress by transcriptional regulation of manganese superoxide dismutase via the SIRT1/FoxO3a pathway. Circulation 127, 1692–701.

Li, W., Jue, T., Edwards, J., Wang, X. and Hintze, T. H. (2004). Changes in NO bioavailability regulate cardiac O2 consumption: control by intramitochondrial SOD2 and intracellular myoglobin. Am J Physiol Heart Circ Physiol 286, H47–54.

Mahaboob Basha, P. and Saumya, S. M. (2013). Suppression of mitochondrial oxidative phosphorylation and TCA enzymes in discrete brain regions of mice exposed to high fluoride: amelioration by Panax ginseng (Ginseng) and Lagerstroemia speciosa (Banaba) extracts. Cell Mol Neurobiol 33, 453–64.

Miar, A., Hevia, D., Munoz-Cimadevilla, H., Astudillo, A., Velasco, J., Sainz, R. M. and Mayo, J. C. (2015). Manganese superoxide dismutase (SOD2/MnSOD)/catalase and SOD2/GPx1 ratios as biomarkers for tumor progression and metastasis in prostate, colon, and lung cancer. Free Radic Biol Med 85, 45–55.

Millay, D. P. and Olson, E. N. (2013). Making muscle or mitochondria by selective splicing of PGC-1alpha. Cell Metab 17, 3–4.

Padmaja Divya, S., Pratheeshkumar, P., Son, Y. O., Vinod Roy, R., Andrew Hitron, J., Kim, D., Dai, J., Wang, L., Asha, P., Huang, B. et al. (2015). Arsenic Induces Insulin Resistance in Mouse Adipocytes and Myotubes Via Oxidative Stress-Regulated Mitochondrial Sirt3-FOXO3a Signaling Pathway. Toxicol Sci 146, 290–300.

Park, S. J., Ahmad, F., Philp, A., Baar, K., Williams, T., Luo, H., Ke, H., Rehmann, H., Taussig, R., Brown, A. L. et al. (2012). Resveratrol ameliorates aging-related metabolic phenotypes by inhibiting cAMP phosphodiesterases. Cell 148, 421–33.

Pi, H., Xu, S., Reiter, R. J., Guo, P., Zhang, L., Li, Y., Li, M., Cao, Z., Tian, L., Xie, J. et al. (2015). SIRT3-SOD2-mROS-dependent autophagy in cadmium-induced hepatotoxicity and salvage by melatonin. Autophagy 11, 1037–51.

Qiu, X., Brown, K., Hirschey, M. D., Verdin, E. and Chen, D. (2010). Calorie restriction reduces oxidative stress by SIRT3-mediated SOD2 activation. Cell Metab 12, 662–7.

Ramis, M. R., Esteban, S., Miralles, A., Tan, D. X. and Reiter, R. J. (2015). Protective Effects of Melatonin and Mitochondria-targeted Antioxidants Against Oxidative Stress: A Review. Curr Med Chem 22, 2690–711.

San-Miguel, B., Crespo, I., Sanchez, D. I., Gonzalez-Fernandez, B., Ortiz de Urbina, J. J., Tunon, M. J. and Gonzalez-Gallego, J. (2015). Melatonin inhibits autophagy and endoplasmic reticulum stress in mice with carbon tetrachloride-induced fibrosis. J Pineal Res 59, 151–62.

Siu, A. W., Maldonado, M., Sanchez-Hidalgo, M., Tan, D. X. and Reiter, R. J (2006). Protective effects of melatonin in experimental free radical-related ocular diseases. J Pineal Res 40, 101–9.

Sundaresan, N. R., Gupta, M., Kim, G., Rajamohan, S. B., Isbatan, A. and Gupta, M. P. (2009). Sirt3 blocks the cardiac hypertrophic response by augmenting Foxo3a-dependent antioxidant defense mechanisms in mice. J Clin Invest 119, 2758–71.

Taghipour, N., Amini, H., Mosaferi, M., Yunesian, M., Pourakbar, M. and Taghipour, H. (2016). National and sub-national drinking water fluoride concentrations and prevalence of fluorosis and of decayed, missed, and filled teeth in Iran from 1990 to 2015: a systematic review. Environ Sci Pollut Res Int 23, 5077–98.

Tao, R., Coleman, M. C., Pennington, J. D., Ozden, O., Park, S. H., Jiang, H., Kim, H. S., Flynn, C. R., Hill, S., Hayes McDonald, W. et al. (2010). Sirt3-mediated deacetylation of evolutionarily conserved lysine 122 regulates MnSOD activity in response to stress. Mol Cell 40, 893–904.

Tseng, A. H., Shieh, S. S. and Wang, D. L. (2013). SIRT3 deacetylates FOXO3 to protect mitochondria against oxidative damage. Free Radic Biol Med 63, 222–34.

Varol, E. and Varol, S. (2012). Effect of fluoride toxicity on cardiovascular systems: role of oxidative stress. Arch Toxicol 86, 1627.

Venegas, C., Garcia, J. A., Escames, G., Ortiz, F., Lopez, A., Doerrier, C., Garcia-Corzo, L., Lopez, L. C., Reiter, R. J. and Acuna-Castroviejo, D. (2012). Extrapineal melatonin: analysis of its subcellular distribution and daily fluctuations. J Pineal Res 52, 217–27.

Wei, L., Zhou, Y., Qiao, C., Ni, T., Li, Z., You, Q., Guo, Q. and Lu, N. (2015). Oroxylin A inhibits glycolysis-dependent proliferation of human breast cancer via promoting SIRT3-mediated SOD2 transcription and HIF1alpha destabilization. Cell Death Dis 6, e1714.

Wu, H., Wu, Y., Ai, Z., Yang, L., Gao, Y., Du, J., Guo, Z. and Zhang, Y. (2014). Vitamin C enhances Nanog expression via activation of the JAK/STAT signaling pathway. Stem Cells 32, 166–76.

Yan, X., Yang, X., Hao, X., Ren, Q., Gao, J., Wang, Y., Chang, N., Qiu, Y. and Song, G. (2015). Sodium Fluoride Induces Apoptosis in H9c2 Cardiomyocytes by Altering Mitochondrial Membrane Potential and Intracellular ROS Level. Biol Trace Elem Res 166, 210–5.

Ye, R., Kong, X., Yang, Q., Zhang, Y., Han, J. and Zhao, G. (2011). Ginsenoside Rd attenuates redox imbalance and improves stroke outcome after focal cerebral ischemia in aged mice. Neuropharmacology 61, 815–24.

Zhou, X., Chen, M., Zeng, X., Yang, J., Deng, H., Yi, L. and Mi, M. T. (2014). Resveratrol regulates mitochondrial reactive oxygen species homeostasis through Sirt3 signaling pathway in human vascular endothelial cells. Cell Death Dis 5, e1576.

Zhou, Y., Qiu, Y., He, J., Chen, X., Ding, Y., Wang, Y. and Liu, X. (2013). The toxicity mechanism of sodium fluoride on fertility in female rats. Food Chem Toxicol 62, 566–72.

